# “All-in-one” Single-Cell Proteomic Analysis of Protein Alterations in Human Oocytes Undergoing in Vitro Aging

**DOI:** 10.64898/2026.01.05.697813

**Authors:** Jue Zhang, Yuting Lu, Shuoping Zhang, Xingyao Wang, Jiao Lei, Feitai Tang, Shen Zhang, Ge Lin

**Affiliations:** Clinical Research Center for Reproduction and Genetics in Hunan Province, Reproductive and Genetic Hospital of CITIC-XIANGYA, 410000, Changsha, China; NHC Key Laboratory of Human Stem and Reproductive Engineering, School of Basic Medical Science, Central South University, 410078, Changsha, China

**Keywords:** Proteomics, Oocyte quality, Oocyte maturation, Embryo quality, Embryo development

## Abstract

**Study question:** What specific proteomic changes and molecular markers associated with in vitro aging occur in human oocytes?

**Summary answer:** Single-cell proteomic analysis revealed the protein dynamics during the in vitro aging of oocytes. Subsequent functional analysis highlighted key biological processes affected in aged oocytes and identified several genes as candidate key factors for oocyte and early embryo quality.

**What is known already:** Ovulated oocytes undergo a time-dependent degradation process if fertilization does not occur within a specific window. This aging process leads to morphological, molecular, and epigenetic changes that can compromise oocyte quality and subsequent embryo development. In previous studies, transcriptome sequencing in humans and mice has revealed serial changes in aging oocytes including oxidative stress, mitochondrial dysfunction, DNA damage, and alterations in cell cycle regulation. These changes can lead to apoptosis, chromosomal abnormalities, and epigenetic modifications, all of which negatively affect oocyte quality. However, research regarding protein level changes and regulatory mechanisms in human oocytes during in vitro aging remains unclear.

**Study design, size, duration:** Utilizing “All-in-one” single-cell proteomics, we investigated the protein expression levels in human oocytes at the germinal vesicle (GV) stage and metaphase II (M II) stage following 24 hours of in-vitro aging. We analyzed four groups of oocytes: fresh GV (n=8), in vitro-aged GV (n=9), fresh M II (n=11), and in vitro-aged M II (n=11). This approach allowed for a detailed comparison of proteomic changes associated with in vitro aging in oocytes at different developmental stages.

**Participants/materials, setting, methods:** Oocytes at GV and MII stage were collected for “All-in-one” single-cell quantitative proteomic respectively. Differentially expressed genes were analyzed using the R package DESeq2 or DEP (filtered with q-value≤0.05 and Foldchange≥1.5). EGSEA package was used to perform pathway analysis.

**Main results and the role of chance:** The proteomic analysis of fresh and in vitro-aged human GV and MII oocytes identified 3,268 proteins. In GV oocytes, compared to fresh samples, 73 proteins were upregulated and 90 were downregulated. Gene Ontology (GO) enrichment analyses revealed these DEPs were involved in key biological processes such as lysosome organization, spindle assembly and oxidative stress response. Gene Set Enrichment Analysis (GSEA) revealed a significant down regulation of genes associated with protein translation initiation and methylation pathways, while those in the apoptosis mitochondrial changes were upregulated. In MII oocytes, 63 proteins were upregulated and 69 were downregulated. GO analysis highlighted their involvement in cytoplasmic translation, ribosome biogenesis and oocyte development. GSEA showed that aged MII oocytes exhibited a downregulation of gene sets in protein translation but an upregulation in membrane fusion pathways compared to fresh oocytes. Furthermore, we identified proteins with consistent expression trends across both GV and MII stages, including MRFAP1 and MT2A, as candidate biomarkers for human oocyte aging in vitro.

**Limitations, reasons for caution:** Single cell proteomic techniques may not fully capture the dynamic range of proteins present within oocytes, leading to potential underrepresentation of low-abundance proteins that could play crucial roles. Furthermore, the heterogeneity within oocytes introduces variability that can obscure the identification of consistent proteomic signatures associated with oocyte quality or aging.

**Wider implications of the findings:** These findings not only provide an understanding of the protein changes underlying the in vitro aging of human oocytes but also offer potential biomarkers and intervention targets for future research and clinical applications in assisted reproductive technologies.

**Study funding/Competing interest(s):** This project received funding from the National Natural Science Foundation of China (NO.82471693, 22574175); the Natural Science Foundation of Hunan Province (NO. 2024JJ4100), the Hunan Provincial Grant for Innovative Province Construction (2019SK4012), the Hundred Youth Talents Program of Hunan Province (to S.Z.), the Major Scientific Program of CITIC Group (No. 2023ZXKYB34100) and Scientific Research Foundation of Reproductive and Genetic Hospital of CITIC-XIANGYA (YNXM-202313, YNXM-202319, YNXM-202211). The authors have no conflicts of interest to declare.

## INTRODUCTION

After gonadotropin surge stimulation, oocytes in prophase I resume meiosis, progressing through germinal vesicle breakdown (GVBD), peripheral spindle migration, and first polar body extrusion (PBE). After these maturation events, oocytes arrested at meiotic metaphase II (MII) can be successfully fertilized if the insemination occurs in a restricted time window after ovulation, within 12h for rodents and 24h for monkeys and humans (Di Nisio et al., 2022). However, oocytes undergo a time-dependent process of degradation, known as “postovulatory aging”. This process is different from “ovarian ageing” which refers to the natural deterioration of ovarian function in females as they near the end of their reproductive lifespan and approach menopause. In vivo, delayed fertilization leads to a series of changes in oocytes, reducing their developmental competence and increasing the risk of failed embryo development. For human, oocytes collected during the assisted reproduction technology (ART) procedures may undergo extended in vitro culture before fertilization, which may lead to in vitro aging (IVA), compromising embryo viability (Lord and Aitken, 2013). In comparison with oocytes analyzed soon after ovulation, these aged oocytes show several morphological, molecular, genomic, and epigenetic abnormalities, which can even lead to apoptosis.

In vivo or in vitro aged oocytes undergo morphological alterations include zona pellucida hardening, cortical granules (CGs) abnormal distribution patterns, CGs vacuolization, and spindle organization (Di Nisio et al., 2022, Lord and Aitken, 2013, Martin et al., 2022). In mouse oocytes, 24 h after ovulation, with the loss of cortical anchorage, more swollen CGs are transported from the membrane into the central region of the oocyte (Zheng et al., 2016). In addition, the decreased numbers of CGs, together with a time-dependent increase in CGs exocytosis, as well as ZP hardening, have been observed in mouse POA-oocytes(Xu et al., 1997). Aged oocytes after ovulation also display aberrant spindle structures, such as elongated, monopolar, multipolar, or disorganized spindles with astral microtubules, as well as multiple cytoplasmic asters (Liang et al., 2018, Li et al., 2024). Notably, in vitro POA oocytes enclosed by cumulus cells (CCs) exhibited partial CGs exocytosis and ZP hardening. But few CC-free oocytes rarely released their CGs, indicating that CCs may accelerate POA progression in vitro (Miao et al., 2005, Kong et al., 2018, Wen et al., 2023). A study on human oocyte demonstrated significant ultrastructural changes including decreased CGs and microvilli, and spindle abnormalities in *in vitro* aged M□ oocyte (Bianchi et al., 2015). Mitochondrial dysfunction is another key factor both in vitro and in vivo oocyte aging. Transmission electron microscopy (TEM) analysis revealed that human IVA oocytes exhibited a decreased number and size of mitochondrial-smooth endoplasmic reticulum (M-SER) aggregates, and an increased amount and size of multivesicular (MV) complexes (Bianchi et al., 2015). Studies have shown that in vitro-aged mouse oocytes exhibit impaired mitochondrial function, characterized by a reduction in mitochondrial membrane potential (MMP) (Zhang et al., 2011) and reduced ATP production, and increased ROS generation (Lord et al., 2015, Martin et al., 2022). Beyond Mt ROS production, lipid peroxidation also serves as a contributing factor that accelerates ROS generation during post-ovulatory oocyte aging (Lord et al., 2015). Lipid aldehydes are formed during lipid peroxidation, a process initiated by ROS-mediated oxidation of polyunsaturated fatty acids (PUFA), where ROS produced via electron leakage from the ETC. But once produced they can then amplify ROS production and contribute to the propagation of further lipid oxidation (Mihalas et al., 2017, Aitken, 2020). Some study reveals that lipid aldehydes including 4-hydroxynonenal (4HNE), malondialdehyde and acrolein are increased during the post-ovulatory MII oocyte aging (Mihalas et al., 2017). Additionally, in both mouse and pig oocytes, alteration in Ca^2+^ oscillation in oocytes are associated with endoplasmic reticulum (ER) Ca^2^□ store depletion, which lead to MMP elevation, triggering pro-apoptotic factor release and ultimately causing POA oocytes fragmentation (Szpila et al., 2019, Yuan et al., 2021).

Both in vivo and in vitro aging have been associated with reduced fertilization rates, poor embryo quality, implantation failure, and abnormalities in the offspring (Takahashi et al., 2013, Martin et al., 2022). Many studies demonstrates reduced fertilization and embryonic development rates of in vitro aged MII oocytes in many species including mice, rats, cows, pigs and humans (Liu et al., 2009, Chian et al., 1992, Kikuchi et al., 2002, Dai et al., 2017, Lord and Aitken, 2013, Takahashi et al., 2013, Nakagata et al., 2024). Certainly, in clinical practice, rescue ICSI may enable the fertilization of post-ovulatory aged oocytes that failed to fertilize 18–24 hours after conventional insemination. But the pregnancy and implantation rates were only about 10% and 5% respectively (Paffoni et al., 2021). Additionally, fertilized in-vitro -aged oocytes exhibit elevated embryonic fragmentation rate (Takahashi et al., 2013, Yamada and Egli, 2017).

Oocyte research is challenging due to the difficulty in sample collection, particularly for human oocytes. However, advances in single-cell omics technologies have provided new insights into the molecular changes occurring during oocyte maturation and early embryonic development (Zhang et al., 2018b, Zhu et al., 2025, Wang et al., 2024). In fact, omics studies have already been conducted on in vitro aged mouse oocytes, revealing key molecular signatures associated with aging (Wang et al., 2017, Guo et al., 2025). But most recent studies on human oocyte aging in vitro have focused on transcriptomic profiling, which cannot accurately reflect protein-level dynamics at the single-cell level. Multiple studies have revealed a significant gap between mRNA and protein expression, highlighting the limitations of relying solely on transcriptomics (Guo et al., 2025, Jiang et al., 2025, Gao et al., 2017). Therefore, single-cell proteomics analysis of human oocytes could offer a deeper understanding of the molecular alterations that occur during in vitro oocyte aging. Despite the existence of some proteomic studies on oocytes, the dynamic proteomic changes associated with oocyte aging in vitro remain poorly understood.

In this study, utilizing a novel “All-in-one” single-cell proteomics method with low-loss sample pre-treatment, we investigated the protein expression levels in human oocytes at the GV and MII stage following 24 hours of in-vitro aging. Our work defines the proteomic differences before and after human oocyte aging in vitro, investigates how dynamic changes during maturation are affected, and aims to identify key regulators or molecular markers responsible for the decline in embryonic quality following oocyte aging.

## MATERIALS AND METHODS

### Human Oocyte collection and culture

This study included 39 human oocytes (8 fresh GV oocytes, 9 IVA GV oocytes, and 11 fresh M□ oocytes and 11 IVA M□ oocytes) from 26 donors. Informed consents were obtained from all donors in this study. This study has been approved by the Ethical Committee of Reproductive and Genetic Hospital of CITIC-XIANGYA (approval No. LL-SC-2024-001). All the human oocyte-related experiments in this study are in compliance with these relevant ethical regulations. The fresh GV and MI oocytes were donated by patients taking intracytoplasmic sperm injection (ICSI) treatments, and these immature oocytes (GV) were not used in regular clinical practice, because the patient already has enough oocytes for their own use. Fresh MII oocytes were from denuded MI oocytes that were kept in in vitro maturation (IVM) medium at 37 °C with 5% CO_2_ for 16-20h. The IVM medium consists of blastocyst medium (Cook Medical, USA) supplemented with 0.075 IU/ml recombinant FSH (Gonal-F; Merck Serono, Germany), 0.1 IU/ml recombinant HCG (Ovidrel; Merck-Serono) and 10% (v/v) of the inactivated patient’s serum (Li et al., 2025). The fresh GV oocytes aged in vitro in medium contain 10μM milrinone (MedChemExpress, HY-14252) for 24h at 37 °C with 5% CO_2_. The fresh MII oocytes aged in vitro in IVM medium for 24h at 37 °C with 5% CO_2_.

### Proteomic Sample Preparation

Single oocyte dissolved in 10□µL of 0.1% DDM in 50□mM Triethylammonium bicarbonate (TEAB) were sonicated at 1-min intervals for 10 times over ice for cell lysis and centrifuged for 3□min at 3000□g. 1□µL of 100□mM DL-dithiothreital (DTT) in 50□mM TEAB was added to the PCR tube. Samples were incubated at 75□°C for 1□h for denaturation and reduction. After that, 2.5□µL of 100□mM Iodoacetamide (IAA) in 50□mM TEAB was added to the PCR tube. Samples were incubated in the dark at room temperature for 30□min for alkylation. 1□µL of 0.05 µg/µL trypsin (Cat# T6567, Sigma) in 50□mM TEAB was added to the PCR tube. Samples were digested for 2h at 37□°C with gentle sharking at ∼350□g.

### LC-MS/MS Analysis

Digested peptides were directly loaded onto a 2 cm PepMap trap column (Thermo Fisher Scientific) for online desalting. Subsequent separation was performed on a 25 cm PepMap analytical column (Thermo Fisher Scientific) using a 102-min gradient from 3% to 38% buffer B (80% ACN, 0.1% formic acid), followed by a 10-min wash at 100% B. FAIMS compensation voltages cycled between -35 V, -45 V, and -65 V with a 1-second duty cycle. MS1 scans (350-1500 m/z) were acquired in the Orbitrap mass analyzer at 60,000 resolution (RF lens: 50%; AGC target: standard; maximum injection time: auto). Dynamic exclusion duration was set to 20 s. MS2 spectra were collected in the linear ion trap in centroid mode (isolation window: 1.6 Th; scan rate: rapid; HCD collision energy: 35%; AGC target: standard; max IT: auto).

### Database Searching

Raw MS data were processed using Proteome Discoverer Software (v2.5, Thermo Fisher Scientific) for feature detection, database searching, and protein/peptide quantification (Peng et al., 2023). The search of MS/MS spectra was conducted against the UniProt human database (downloaded on June 27th, 2022, containing 79,435 entries). Carbamidomethylation on cysteine residues was set as fixed modification. Methionine oxidation and N-terminal protein acetylation were chosen as variable modifications. The mass tolerance for precursors and fragments were set to 10 ppm and 0.6 Da, respectively. Minimum and maximum peptide lengths were six and 144 amino acids, respectively. The maximum missed cleavage for every peptide was two. Protein identifications were filtered at a 1% false discovery rate (FDR). Remaining parameters employed Proteome Discoverer defaults.

### Data Analysis

To ensure the stability of MS-quantified proteins within experimental groups, we filtered out proteins detected in fewer than three samples across all groups. The imputation was processed by DEP (version 1.28.0, R package) (Zhang et al., 2018a). Proteins differences between groups was tested by limma (version 3.62.2, R package) (Ritchie et al., 2015). and the significantly differential proteins were filtered by p ≤ 0.05 & fold change ≥ 1.5.)GO functional enrichment analysis and GSEA was performed using clusterProfiler (version 4.14.6, R package), and the correspondent gene sets selected here were retrieved from the C5 ontology gene sets which were downloaded from https://www.gsea-msigdb.org/gsea/msigdb/index.jsp.

### Mice strain, superovulation, and fertilization

Wild-type C57BL/6 mice were obtained from the China Hunan Slac Jingda Laboratory Animal Co., Ltd. Mice were maintained under specific pathogen free conditions in a controlled environment of 20–22 °C, with a 12/12 h light/dark cycle, 50–70% humidity, and food and water provided. For superovulation and fertilization, 4-week-old female mice were injected with 5 IU of pregnant mare serum gonadotropin (PMSG; Hangzhou animal pharmaceutical). After 44 h, the mice were injected with 5 IU human chorionic gonadotropin (hCG; Hangzhou animal pharmaceutical) and mated with adult males. Successful mating was confirmed based on the presence of a vaginal plug. All procedures in this study were conducted in accordance with the Guide for the Care and Use of Laboratory Animals and were approved by the Animal Welfare Ethics Committee of Central South University (approval no. XMSB-2024-0466).

### Zygote collection, in vitro culture and treatment with inhibitor

Mouse zygotes were obtained from oviducts by inducing superovulation in 4-week-old females that were mated with males of the same strain. Zygotes were released in 37 °C pre-warmed M2 medium (Sigma-Aldrich, M7167) and cultured in mini-drops of M16 medium (Nanjing Aibei Biotechnology, M1735) covered with mineral oil (Nanjing Aibei Biotechnology, M2460) at 37 °C in a 5% CO_2_ atmosphere. Embryos that reached the 2-cell, 4-cell, 8-cell, morula, and blastocyst stages were obtained at 36, 48, 60, 72, and 84 h post-hCG injection. During in vitro embryo culture, we transported the embryos into the fresh culture medium every 48 h. For FSCN inhibition, zygotes were treated with inhibitor diluted in M16 medium covered with mineral oil. The inhibitors used are listed in Supplementary Table 8.

### Human 3pn embryo collection, in vitro culture and treatment with inhibitor

All of the embryos were obtained with signed informed consent by the donor couples. In vitro fertilized 3pn embryos were cultured until the 8-cell stage using a G-1 plus (Vitrolife) human embryos culture medium. For inhibitor treatment, 3pn embryos were cultured with FSCN inhibitor diluted in G-1 plus medium. The inhibitors used are listed in Supplementary Table 8.

### Immunofluorescent staining

Embryos were fixed with 4% paraformaldehyde in phosphate-buffered saline (PBS) at room temperature for 20 min and then permeabilized with 0.3% Triton X-100 in PBS for one night at 4°C. After blocking with 1% BSA (bovine serum albumin) in PBST, embryos were incubated with primary antibodies diluted in blocking solution at room temperature for 2 h, then incubated with Alexa Fluor 594-conjugated or 488-conjugated secondary antibodies at room temperature for 2 hrs. Imaging of embryos after immunofluorescence was performed on a Zeiss CellDiscover7 confocal microscope. The antibodies used are listed in Supplementary Table 8.

## RESULTS

### Optimization of an “All-in-One” Workflow for Single Human Oocyte Proteomics

To enhance proteome coverage in single-oocyte analysis, we developed and optimized a n-Dodecyl β-D-maltoside (DDM)-assisted “All-in-one” sample preparation workflow, building upon our prior methodology (Guo et al., 2023). This integrated approach consolidates cell lysis, protein denaturation, reduction, alkylation, and digestion within a single 200 µL PCR tube. Optimization in cell lysis and trypsin digestion was performed using 500 HEK293T cells. The reason why choosing 500 HEK293T cells while not single human oocyte directly for optimization was because: firstly, the protein amount of a single human oocyte is comparable to ∼500 somatic cells (Virant-Klun et al., 2016), and secondly, the inherent oocyte heterogeneity complicates attributing differences in results solely to methodological variations.

To optimize cell lysis condition, different DDM concentration ranging from 0.01% to 0.2% was tested. Lysis with 0.1% DDM yielded the highest number of protein and peptide identifications (Fig. S1A, B). Proteins identified with 0.1% DDM spanned a dynamic range of 4.87 orders of magnitude, larger than that with lower DDM concentration (Fig. S1C). Although 0.2% DDM achieved a wider dynamic range (5.25 orders), identification numbers significantly decreased, likely due to LC column saturation by higher DDM concentration impairing chromatographic performance. The log2 intensity of the peptides commonly identified across different conditions was higher with 0.1% DDM (Fig. S1D), indicating better protein extraction efficiency. The quantified protein abundance variations among triplicate analysis were lower with 0.1% DDM (Fig. S1E), suggesting better extraction reproducibility. Furthermore, no significant biases in cellular component distribution (Fig. S1F) (Supplemental Table S5), Grand average of hydropathicity (GRAVY) value (Fig. S1G), isoelectric point (pI; Fig. S1H), and molecular weight (MW; Fig. S1I) were observed for proteins identified using 0.1% DDM.

To optimize trypsin digestion, three aspects were evaluated. The first aspect is trypsin source. Four commercially available trypsins were compared. Sigma, Hualishi, and Promega trypsins yielded significantly more protein and peptide identifications than Thermo trypsin (Fig. S2A, B). Sigma trypsin provided the lowest median quantification coefficient of variation (CV), indicating better digestion consistency and reproducibility (Fig. S2C), thus was selected for subsequent analysis. The second aspect is trypsin-to-substrate ratio (w/w). Ratios from 100:1 to 1:1 were tested. A ratio of 10:1 achieved optimal protein and peptide identification (Fig. S2D, E). Although a 50:1 ratio offered slightly better digestion consistency (Fig. S2F), the significant identification advantage of 10:1 led to its selection. The third aspect is digestion time. Shorter digestion times yielded higher identification numbers (Fig. S2G, H), potentially due to under a high trypsin-to-substrate ratio, prolonged digestion time leads to autolysis of trypsin and generation of excessively short peptides, both of which are detrimental to mass spectrometry detection (Yuan et al., 2017). While longer time improved quantification reproducibility (Fig. S2I), the identification advantage and higher throughput of a 2-hour digestion warranted its adoption.

In order to accurately access the effect of different mass spectrometry (MS) parameters on the analysis results, the variations in sample preparation steps need to be avoided and thus 100 ng K562 digest was used in the optimization of MS parameters (Wu et al., 2023). Key instrument parameters including compensation voltage (CV), MS resolution, MS/MS auto gain control (AGC) target, MS/MS maximum injection time and MS/MS acquisition number (Fig. S3A) were evaluated for their impact on identification and quantification reproducibility. Method 3 yielded significantly more protein and peptide identifications compared to other methods (Fig. S3B, C). Its identified proteins spanned 5.4 orders of magnitude, exceeding the dynamic range achieved by other methods (Fig. S3D). Peptides commonly identified across different methods showed higher log2 intensities with Method 3 (Fig. S3E), demonstrating enhanced MS sensitivity. Although the median CV for Method 3 was slightly higher than those of other methods (Fig. S3F), its significant advantages in identification and sensitivity led to its selection in our “All-in-one” single cell proteomics workflow.

Finally, the optimized “All-in-One” single cell proteomics workflow was tested using single human oocytes, and the average number of proteins quantified per oocyte in GV and MI were 3185±167(n=6) and 3283±49 (n=3) (Fig. S4A).

### “All-in-one” Single-Cell Quantitative Proteomic Profiling of Human in-vitro-Aging Oocyte

To profile human oocyte in vitro aging, we used 39 human oocytes (8 fresh GV oocytes, 9 IVA GV oocytes, and 11 fresh M□ oocytes and 11 IVA M□ oocytes) from 26 donors (Supplemental Table S1). On the day of oocyte retrieval, immature oocytes, including GV and MI stage oocytes, are collected. Fresh GV oocytes are directly sampled immediately after retrieval. MI oocytes, on the other hand, are subjected to in vitro culture until they reach the MII stage. For studies involving aged oocytes, these GV and MII oocytes are further cultured for an additional 24 hours to induce in vitro aging (Fig. 1A). This process allows for the collection and analysis of oocytes at different stages of maturation and aging. The typical morphology of collected oocytes was as shown in Fig1B. Each oocyte was subjected to cell lysis, protein extraction, denaturation, reduction, alkylation, trypsin digestion and LC-MS/MS analysis (Fig. 1A). We filtered out proteins detected in fewer than three samples across all groups. Totally, 3268 proteins and 29,735 peptides were identified (Supplemental Table S2). The average number of proteins/peptides quantified per oocyte in fresh or IVA GV, and MII stage were 2830/16173, 2768/15825, 2804/15898 and 2793/16248, respectively (Fig. 1C, S4B-C). To assess data quality, we compared our results with Xie et al.’s single-cell RNA-seq and ribo-seq data, identifying 24,861 expressed genes in single oocyte. Our data matched 3,057 proteins (93.54%) (Fig. S4C). The protein levels increased progressively with prolonged in vitro culture time, as indicated by the total abundance of each group (Fig.1E). Additionally, compared to Guo et al.’s single-cell proteomics data which identified 2434 proteins in single oocyte, 69.26% (1686/2434) of the proteins were also identified in our dataset (Fig. S4D). There are 1052 proteins detected in our data are maternal genes (Fig. 1D, Fig. S4F-G). The Biological processes functional enrichment of these maternal proteins is shown in Figure 1F, which includes critical biological processes in oocytes such as carboxylic acid metabolic process, vesicle-mediated transport, mitochondrion organization, and citrate cycle (TCA cycle).

**Figure 1.**
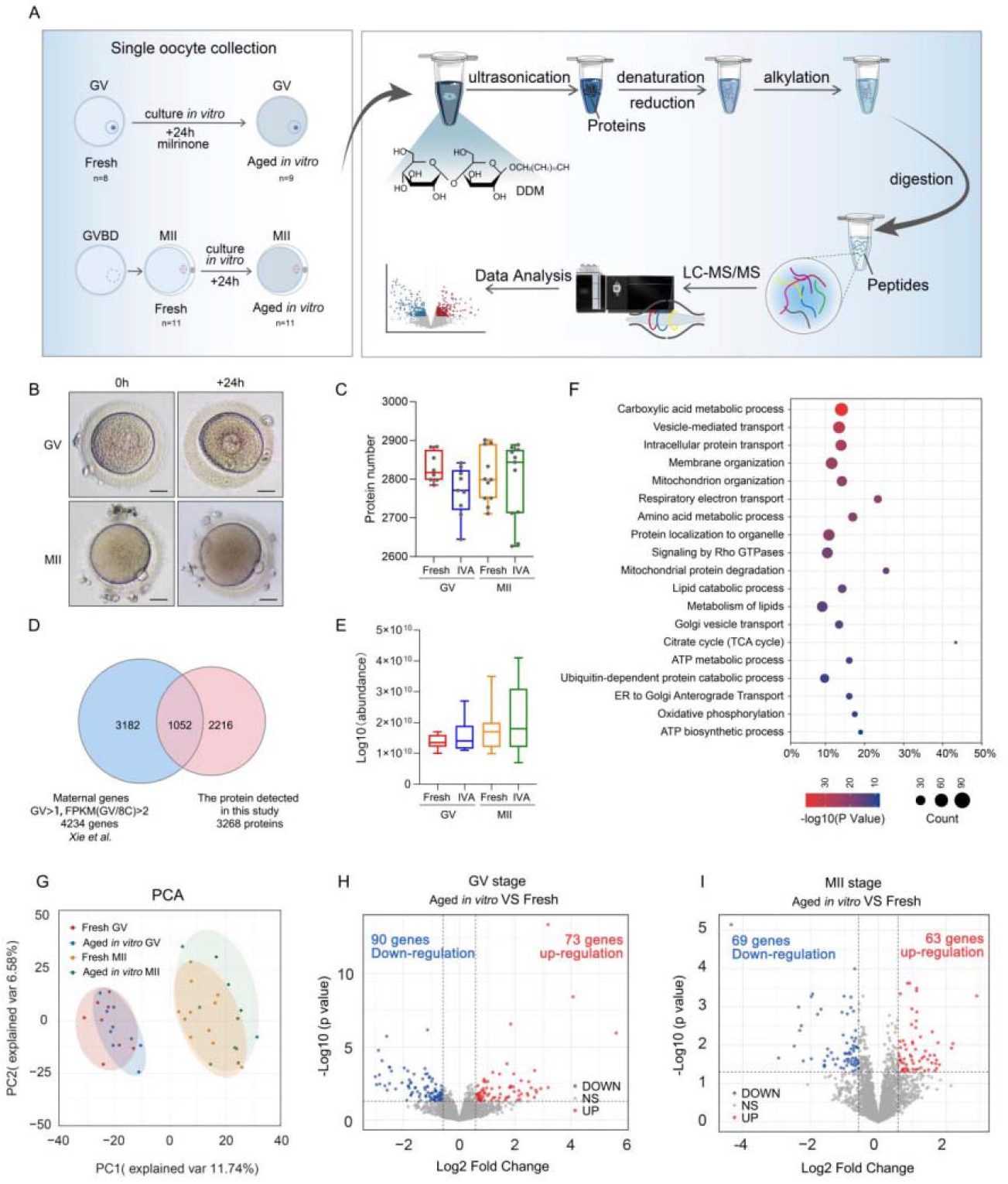
Single-cell quantitative proteomic profiling of human oocytes aging in vitro. A. schematic illustration of single-cell quantitative proteomic profiling of human oocyte aging in vitro. B. DIC representative images of GV, M□ stage oocytes that collected in fresh or aged in vitro 24h condition. The scale bar represents 25 μm. C. The box plot illustrates the number of proteins detected by “All-in-one” single cell proteomics method for each oocyte. D. Venn diagram showing the overlap between the identified proteins in single-cell proteomics dataset (this study) and maternal genes (FPKM≥1 in GV stage) in single cell RNA-seq (Xie et al). E. The box plot displays the total abundance of protein expression for each group, with the data presented on a log10 scale. F. The enriched biological processes analysis of the overlap genes in D. G. Principal component analysis (PCA) of the proteomics data from 39 fresh or in-vitro-aged (IVA) human oocytes including GV and M□ stage. H. Volcano plot shows the different expression proteins (DEPs) between the IVA and fresh group in GV oocytes. Fold change ≥1.5, p-value ≤ 0.05. I. Volcano plot shows the protein expression differences between the IVA and fresh group in M□ oocytes. Fold change ≥ 1.5, p-value ≤ 0.05.

We performed PCA analysis of the fresh and in-vitro aging oocytes (Fig.1G). Based on our proteomic data, oocytes from different stages separated into two clusters. GV oocytes exhibited relatively compact distribution, while MII oocytes showed more dispersed. Reproducibility analysis (Fig. S4H) of protein quantification among the oocyte samples indicated high reproducibility, with pairwise Pearson’s correlation coefficients ranging from 0.89 to 0.94 and 0.86 to 0.94 among oocytes in GV and M□ stage. Oocytes from the same donor showed slightly greater similarity to each other with relatively higher Pearson’s correlation coefficients from 0.89 to 0.94. (Fig. S4H).

### Comparative Proteomic Profiling Reveals Functional Enrichment in GV and MII Oocytes Undergoing in Vitro Aging

To investigate the effects of in vitro aging on the proteomic profiles of oocytes, we performed differential protein expression analysis in oocyte at GV and MII stage. In GV oocytes, compared to fresh samples, we observed the up regulation of 73 proteins and the down regulation of 90 proteins (Fig. 1H). In MII oocytes, we identified a total of 176 differential proteins (Supplemental Table S3), including 63 upregulated proteins and 69 downregulated proteins (Fig. 1I). The DEPs were classified according to their expression profiles across follicular developmental stages and their differential expression patterns were illustrated in heatmaps (Fig. A-B) (Supplemental Table S4). Stage-specific biological processes enriched in DEPs between fresh and in vitro aged (IVA) GV oocytes across clusters □ to □ (Fig. C-D) (Supplemental Table S5).

After 24 hours of in vitro aging of GV oocytes, most DEPs (Fig.2A) were specific-highly expression during the primary follicular development stage (Cluster □, 49 proteins), and biological process annotation showed enrichment of “regulation of RNA splicing” (9 proteins) (Fig. 2B) (Supplemental Table S5), represented by NCL, the major nucleolar protein of growing eukaryotic cells, that plays a role in pre-rRNA transcription and ribosome assembly (Ugrinova et al., 2007, Yang et al., 2024a) (Fig. 2E). The terms including “actin filament organization” and “actin cytoskeleton organization” (Fig. 2E), as well as the term “organelle localization” (7 proteins), represented by BUB3 were enriched. In cluster □, the term “monocarboxylic acid metabolic process” was significantly enriched with lactate dehydrogenase B (LDHB) identified as a key enzymatic component interconverts lactate/pyruvate and NAD+/NADH (Sharpley et al., 2021). Additionally, the terms “lipid transport” in cluster I and “positive regulation of lipid metabolic process” in cluster □ were also enriched. The representative genes APOA4 (apolipoprotein A4) and APOA2 (apolipoprotein A2) were both upregulated during IVA (Fig. 2E).

**Figure 2.**
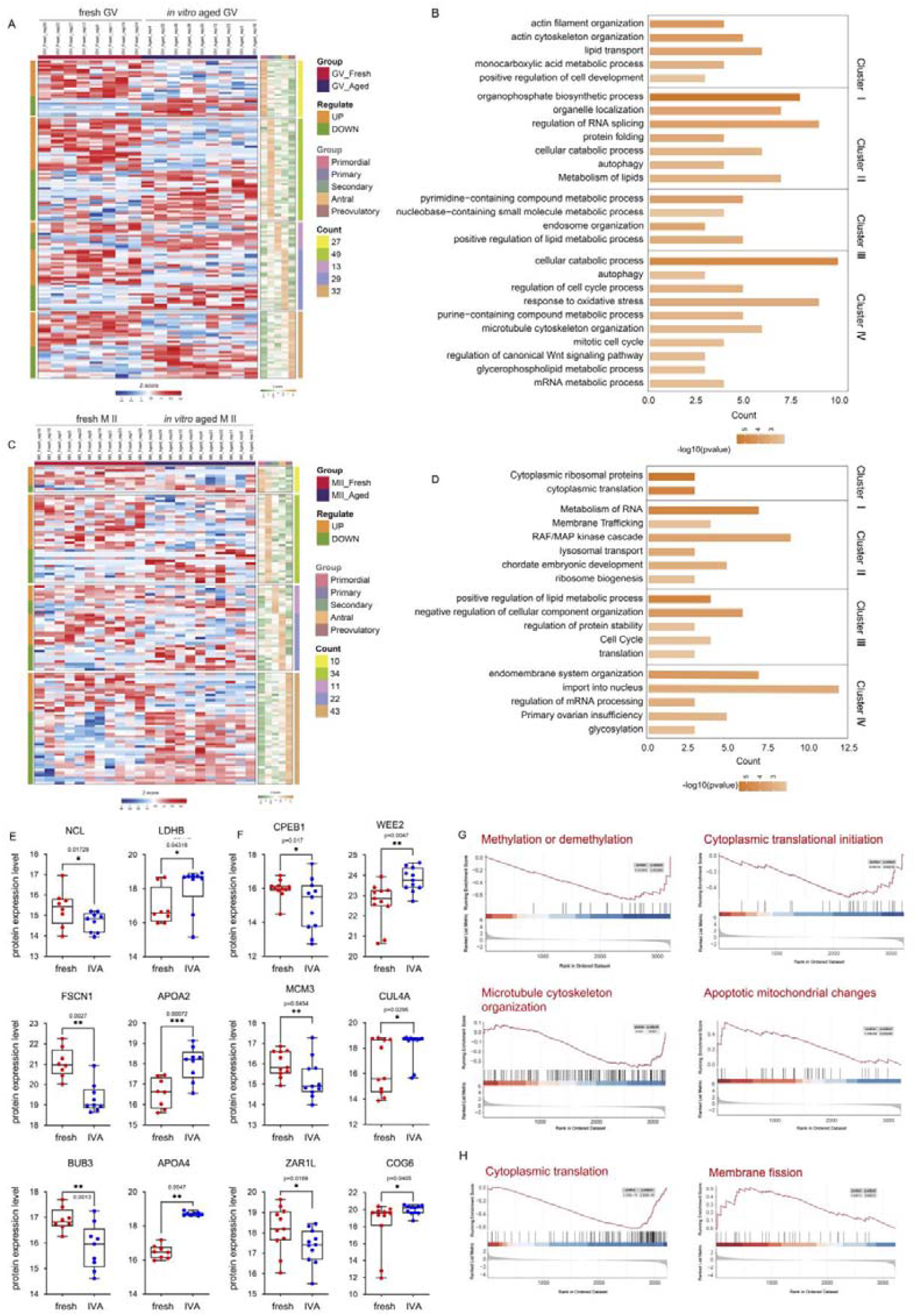
Heatmap and functional enrichment analysis of differential expression protein in human in-vitro-aging oocytes. **A**. Heatmap showing the expression levels of 163 differentially expressed proteins (DEPs) during folliculogenesis (data from GSE107746) and fresh or IVA human GV oocytes (this date). **B**. Stage-specific biological processes enriched in DEPs between fresh and in vitro aged (IVA) GV oocytes across clusters: C □: primordial follicles; C □: primary follicles; C □: secondary and Antral follicles; C □: pre-ovulatory follicles. **C**. Heatmap showing the expression levels of 132 DEPs during folliculogenesis (data from GSE107746) and fresh or IVA human M□ oocytes (this date). **D**. Stage-specific biological processes enriched in DEPs between fresh and in vitro aged (IVA) M D oocytes across clusters: C □: primordial follicles; C □: primary follicles; C □: secondary and Antral follicles; C □: pre-ovulatory follicles. **E**. Box plots show expression of representative proteins selected from selected BP enrichment pathways in GV oocyte, as interleaved boxes and whiskers with minimum and maximum values displayed. * p ≤ 0.05, ** p ≤ 0.01. **F**. Box plots show expression of representative proteins selected from selected BP enrichment pathways in MII oocyte, as interleaved boxes and whiskers with minimum and maximum values displayed. * p ≤0.05, ** p ≤ 0.01. **G**. GSEA enrichment plots of GO pathways between fresh and IVA GV oocytes. **H**. GSEA enrichment plots of GO pathways between fresh and IVA MII oocytes.

In M□ stage oocytes, most DEPs were specific-highly expression during the preovulatory follicular development stage (Cluster □, 43 proteins) (Fig2A). Biological process enrichment analysis revealed DEPs are involved in “import into nucleus”, contained key maternal gene WEE2 (Hanna et al., 2010, Nozawa et al., 2023, Liu et al., 2024b) that increased in MII in vitro aged oocytes ((Fig 2F), and “primary ovarian insufficiency” that involved CPEB1 and CNOT8. In Cluster □, The DEPs were highly enriched in “ribosome biogenesis”, represented by CUL4A. Additionally, this cluster also showed strong enrichment in “cell cycle” including MCM3 and ANKLE2, as well as in translation pathways, particularly involving ZAR1L (Fig. 2F). In Cluster □, the term “endomembrane system organization” was enriched including COG6, one of the subunit of conserved oligomeric Golgi (COG) complex (Rui et al., 2020).

Gene Set Enrichment Analysis (GSEA) revealed significant downregulation of genes associated with “cytoplasmic translation initiation” and “methylation or demethylation” pathways in GV-stage aged oocytes, suggesting that these pathways may be suppressed during in vitro aging. Conversely, upregulation of genes associated with “microtubule cytoskeleton organization” and “apoptotic mitochondrial changes” pathways (Fig.2E). Additionally, in vitro-aged MII oocytes exhibited downregulation of genes involved in “cytoplasmic translation” and upregulation of genes in “membrane fission” compared to fresh oocytes (Fig.2G). The functional enrichment analysis of the DEGs validated the molecular alterations reported in previous studies of in vitro oocyte aging (Di Nisio et al., 2022, Martin et al., 2022).

### Protein Dynamics During in Vitro Aging Oocyte and Its Relationship with Polyadenylation in Oocyte Maturation

During oocyte maturation, a large amount of protein translational activation occurs (Zhang et al., 2020, Liu et al., 2023, Liu et al., 2024a). Based on their expression trends in fresh human oocytes from the GV to MII stage, we categorized the 3268 detected genes into three modes of translational dynamics (Fig.3A) (Supplemental Table S3). The “translation activation” proteins are increased from GV to M□ (Fig.3A), which indicated these genes may play critical roles in oocyte maturation or preparing for post-fertilization embryonic development. Among these proteins, 24 were down-regulated in in vitro aged human MII oocytes (Fig.3B) such as MTO1 and NDUFAF3 (Fig3D). The “maternal clearance” proteins are decreased from GV to M□ (Fig3A). These proteins targeted for degradation during oocyte maturation and also included critical maternal factors and 13 proteins exhibited impaired protein decay in in vitro -aged MII oocytes (Fig.3C). For example, MTOR (a master regulator of translation and autophagy) and EIF3A (a core initiation factor) were impaired decay in IVA M□ oocytes (Fig3D). All proteins displaying aberrant expression trends (37 proteins) are presented in Figure S5A.

Due to transcriptional silencing in oocyte maturation, protein expression regulation relies on post-transcriptional regulation of maternal mRNA and is associated with mRNA poly(A) tail length. Additionally, many maternal mRNA undergo poly(A) tail elongation during GV to MII stages. Therefore, we hypothesized that changes in the protein profile of oocytes subjected to prolonged in vitro culture might be related to their poly(A) tail lengths. To investigate this, we integrated data from the published dataset (HRA003115) (Liu et al., 2024a) and calculated the average poly (A) tail length of DEPs during in vitro aging in GV and MII stage separately (Fig.3E). We find that there was no significant difference in poly (A) tail length between DEPs at GV stage oocytes, while in MII oocytes, proteins exhibiting up-regulated protein during in vitro aging underwent poly(A) tail elongation throughout oocyte maturation, and show longer poly(A) tails than those with down-regulated proteins at M□ stage (Fig3E). Represented DEPs WEE2 and SUN1 both exhibit poly(A) elongation during oocyte maturation and increased after in vitro aged (Fig. 3F-G, J). Conversely, CPEB1 undergoes poly(A) tail shortening and YBX2 exhibits no significant change in poly(A) length. While the protein levels of both declines in in vitro aged M□ oocytes (Fig. 3H-J). Furthermore, extended analysis revealed that among up-regulated proteins at the M□ stage, 48.21% underwent poly(A) tail elongation, while only 14.29% exhibited shortening, with the remainder (37.50%) showing no change. Conversely, among down-regulated proteins, 50.88% maintained stable poly(A) tail lengths, 22.80% displayed shortening, and merely 26.32% underwent elongation (Fig S5B-C). This indicates that some proteins with elongated poly(A) tails during oocyte maturation sustain persistent translation during prolonged in vitro culture, resulting in increased protein abundance.

**Figure 3.**
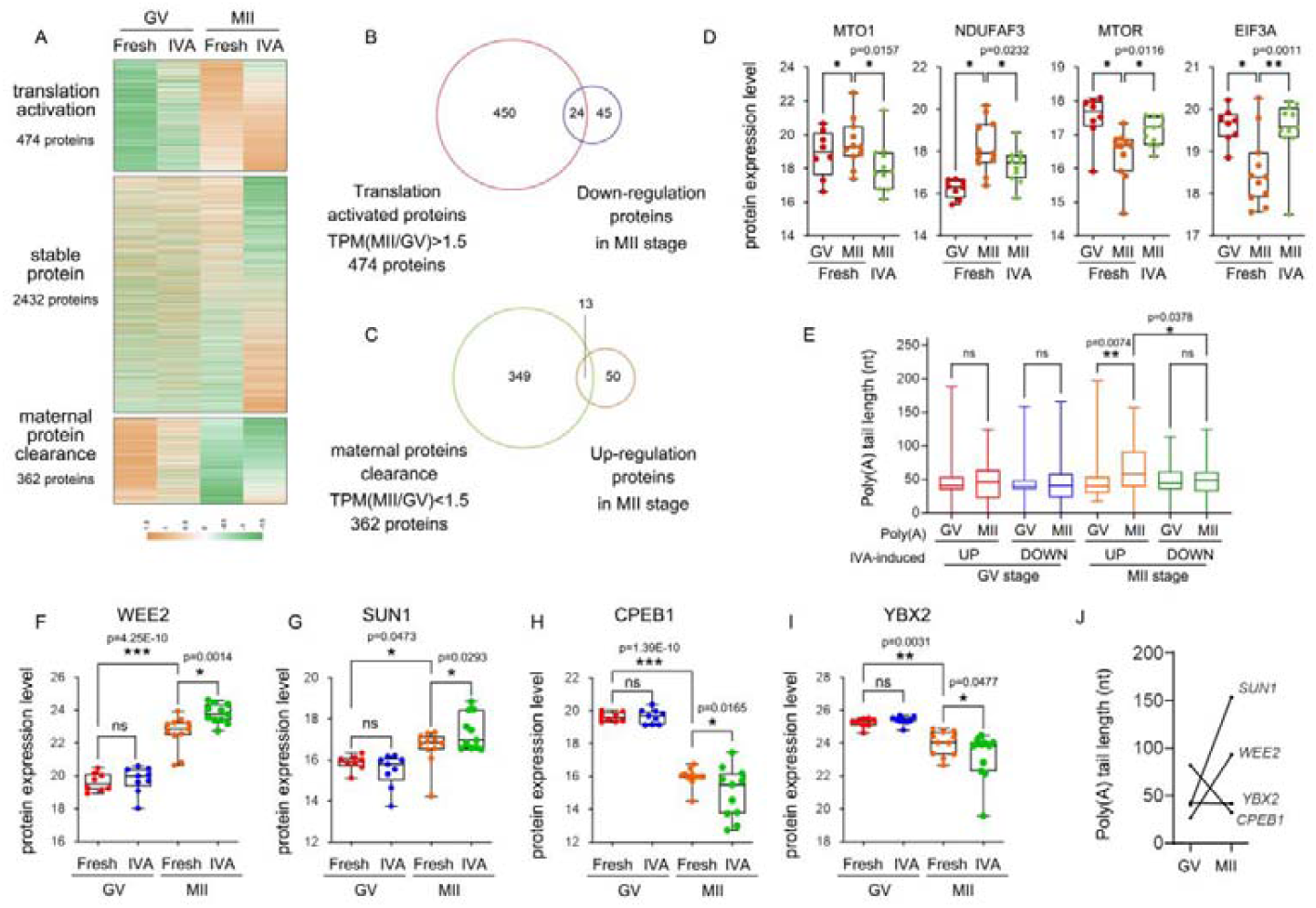
The relationship between protein dynamics in vitro aging and polyadenylation during oocyte maturation. **A**. Heatmaps showing the dynamics of these 3268 identified proteins during oocyte maturation. The translational activated proteins (474) were identified by a fold change ≥1.5 in expression between fresh MII and GV oocytes. The maternal decay proteins (362) identified by a fold change ≥1.5 in expression between fresh GV and M□ oocytes. The remaining proteins (2432) are defined as stably expressed proteins from GV to MII stage. **B**. Venn diagram showing the overlap between translational activated proteins and 69 down regulated proteins in IVA M □ oocytes. **C**. Venn diagram showing the overlap between maternal decay proteins and 63 up regulated proteins in IVA M □ oocytes. **D**. Box plots show expression of representative proteins selected from the proteins with aberrant expression patterns. * p ≤ 0.05, ** p ≤ 0.01. **E**. Box plots show the mean poly(A) tail length of DEPs after in vitro aging during oocyte maturation. * p ≤ 0.05, ** p ≤ 0.01. **F-I**. Box plots show expression of representative proteins. * p ≤ 0.05, ** p ≤ 0.01, *** p ≤ 0.001. **J**. The poly(A) tail length of representative proteins during oocyte maturation.

## DISCUSSION

In clinical assisted reproductive technology (ART), prolonged in vitro culture or delayed fertilization (e.g., rescue ICSI) can subject oocytes to post-maturation aging, compromising fertilization potential and subsequent embryo development. While previous studies have characterized morphological, organelle dynamics, and Ca^2^□ homeostasis in POA oocytes, our study provides the first comprehensive single-cell proteomic profiling of human oocytes undergoing 24-hour in vitro aging at both GV and MII stages. This approach delineates the functional impact of in vitro aging on oocyte maturation and identifies novel molecular biomarkers for oocyte quality assessment and potential therapy targets for improving aging oocytes. Furthermore, it may also provide a viable model system for investigating the mechanistic insights into fragmentation in early human embryo development.

In this work, we developed an easy-to-use, DDM-assisted “All-in-one” workflow for in-depth and reproducible single-cell proteomic analysis. Through systematic optimization of cell lysis, protein extraction, trypsin digestion, and MS detection, this method achieves superior identification depth for single human oocytes compared to other recent in-tube sample preparation studies (Guo et al., 2022, Zhang et al., 2025). Furthermore, its performance is even comparable to results obtained using a chip-based sample preparation method (Dang et al., 2023). While chip-based approaches enhance sample recovery and thereby improve proteomic identification depth, they require significant operator expertise and face inherent technical barriers in chip fabrication, limiting their widespread adoption. Given its excellent performance and operational simplicity, we anticipate this method will serve as a viable and promising technical option for single-cell proteomic studies, particularly those involving single oocytes or embryos.

Our quantitative proteomic analysis revealed that the proportion of DEPs during in vitro aging was relatively small compared to the total proteins detected: 163/3267 (5.0%) at GV and 132/3064 (4.3%) at MII stage. This observation was further supported by PCA result (Fig1G), which showed limited separation between aged and fresh oocytes within the same stage. Collectively, these findings suggest that the oocyte proteome remains relatively stable during the 24-hour in vitro aging period examined. This proteomic stability implies that the functional decline associated with aging might arise from alterations in a critical subset of proteins, potentially offering promising targets for targeted interventions to improve oocyte quality. However, we acknowledge that the limited detection depth inherent to this single-cell proteomic technologies could contribute to this observation, as low-abundance proteins crucial for oocyte function might evade detection. Future studies employing more sensitive methods or targeted approaches could uncover additional relevant changes.

The GO enrichment analysis (Fig2B, D) and GSEA (Fig2 G, H) revealed significant functional alterations in several key areas. Notably, we observed enrichment of BP related to “metabolism of lipids” and “lipid transport” in GV-stage oocytes, “positive regulation of lipid metabolic process” in M□-stage oocytes. Representative protein ACOX1 (Peroxisomal acyl-coenzyme A oxidase 1) was upregulated in aged GV oocytes (Table S5). ACOX1 involved in the initial and rate-limiting step of peroxisomal beta-oxidation of straight-chain saturated and unsaturated very-long-chain fatty acids, producing hydrogen peroxide (H□O□)(Ferdinandusse et al., 2007). This H□O□ may contribute to lipid peroxidation. As lipid peroxidation level are known to be elevated within post-ovulatory aged oocytes (Takahashi et al., 2003), increasing ROS level (Mihalas et al., 2018). Secretory protein APOA2 was increased both in aged GV and MII oocytes (Fig 2 E). APOA2 stabilize HDL (high density lipoprotein) structure by its association with lipids, and affect the HDL metabolism (Sarkar et al., 2024). Previous study found out that protein level of ApoA1 (hub protein of HDL) was increased in follicular fluid of non-pregnant patients (Kurdi et al., 2023). Our data also confirmed significant alterations in proteins associated with “apoptotic mitochondrial changes” and “response to oxidative stress” in GV stage (Fig 2 B, G). These findings corroborate that elevated levels of lipid peroxides, alongside alterations in lipid transport and lipoprotein homeostasis, are prominent features of in vitro aged oocytes. The resulting oxidative stress and changes in the fluidity of cellular membranes and membrane-bound organelles are likely key mechanisms contributing to the decline in quality observed in in vitro aged oocytes.

Furthermore, we identified a specific and notable decrease in the abundance of BUB3 (Fig. 2 E), a core component of the spindle assembly checkpoint (SAC) in MII stage oocytes. SAC ensures that chromosome segregation takes place only after correct end-on kinetochore microtubule attachments to both poles have been achieved (Homer et al., 2005, Li et al., 2009). This finding mirrors previous reports of reduced *Mad2* (another essential SAC protein) mRNA levels in POA mouse oocytes (Steuerwald et al., 2005). This compromised SAC function poses a significant threat to chromosome segregation fidelity, increasing the risk of aneuploidy and contributing to the reduced developmental potential of aged oocytes. In contrast to previous reports of SIRT1 downregulation in POA oocytes (Xing et al., 2021, Zhang et al., 2016), SIRT1 was not detected in our proteomic dataset. We did detect SIRT3, but no significant difference was observed.

Additionally, our analysis of poly(A) tail length dynamics among differentially expressed proteins revealed that select mRNAs undergoing poly(A) tail elongation during oocyte maturation exhibit sustained translational activity throughout prolonged in vitro culture (Fig3E). This persistent translation culminates in the progressive accumulation of corresponding proteins. The maintenance of this translational function hints that the poly(A) tail length-dependent protein translation mechanism remains operational during in vitro aging. However, it remains experimentally unconfirmed whether the increase in such proteins drives oocyte aging or serves as a protective response. This question warrants further investigation in subsequent studies.

Of course, we also aim to utilize these dynamic changes to identify biomarkers of oocyte quality. By focusing on genes with consistent expression trends in both GV and MII stages, we identified some candidate biomarkers for in vitro aged human oocytes, such as APOA2 and MT1H. There is very little research on the functions of these two proteins in oocytes and early embryos. APOA2 (Apolipoprotein A-II) is one of the major constituents of high-density lipoprotein (HDL) and plays a critical role in lipid metabolism and reverse cholesterol transport. HDL is the major lipoproteins detected in substantial amounts in follicular fluid (FF), and FF HDL cholesterol concentration and particle size were negatively correlated with IVF outcomes and preimplantation embryo quality parameters, including fragmentation and symmetry(Browne et al., 2009). APOA2 up-regulated in age oocyte may reflect altered intracellular cholesterol homeostasis. Metallothioneins (MTs) are a group of evolutionarily conserved cysteine-rich metal binding proteins that actively participate in the cellular defense against free radicals. Cysteine residues from Metallothioneins serve to neutralize harmful oxidant radicals (Han et al., 2013). MT1H (Metallothionein-1H) increasing may represent a stress-protective mechanism in oocytes during in vitro aging.

In summary, in vitro aging of oocytes is an issue that cannot be completely avoided in human ART, particularly in situations where extended periods of in vitro maturation are required prior to in vitro fertilization (IVF), or in cases such as rescue ICSI. According to the findings of this study, in vitro aging does exert certain effects on oocytes at the protein level, with several critical pathways and factors being impaired. Therefore, in clinical practice, it is essential to strike a balance—ensuring that the culture duration is sufficiently long to oocyte maturation, yet not excessively prolonged so as to trigger in vitro aging. Beside, despite significant advances in recent years, current single-cell proteomics methods remain fundamentally limited in comprehensively capturing the proteome, especially low-abundance proteins (e.g., transcription factors, signaling molecules) and proteins with potentially extensive post-translational modifications, such as SIRT1. This stems from technical constraints: minimal sample input leads to significant stochastic losses during preparation, while mass spectrometry faces inherent sensitivity and dynamic range boundaries. Therefore, further enhancing the depth and sensitivity of proteomic technologies remains imperative to fully elucidate the molecular mechanisms underlying oocyte aging and to identify more precise biomarkers.

## Data availability

All MS data were deposited in the ProteomeXchange Consortium (https://proteomecentral.proteomexchange.org) via the PRIDE partner repository with identifier PXD057953 and PXD058630. The poly(A) tail length measurements for human oocyte mRNAs were obtained from the publicly available dataset HRA003115.

## Acknowledgements/Funding

The authors wish to thank Prof. Keliang Wu and his team for providing the standardized human oocyte mRNA poly(A) tail length data from the publicly available dataset HRA003115. The authors acknowledge the financial support for this project from the following funding agencies: the National Natural Science Foundation of China (Grants No. 82471693 and 22574175); the Natural Science Foundation of Hunan Province (Grant No. 2024JJ4100); the Hunan Provincial Grant for Innovative Province Construction (Grant No. 2019SK4012); the Hundred Youth Talents Program of Hunan Province (awarded to S.Z.); the Major Scientific Program of CITIC Group (Grant No. 2023ZXKYB34100); and the Scientific Research Foundation of Reproductive and Genetic Hospital of CITIC-XIANGYA (Grants No. YNXM-202313, YNXM-202319, and YNXM-202211).

## Contribution

G.L., S.Z. and J.Z. contributed to conceptualization and experimental design. J.Z. and S.Z. contributed to experimental design, data curation and manuscript writing. Y.L. and S.Z. were responsible for oocyte collection.

X.W. performed formal analysis and data curation. J.L. and F.T. contributed to methodology, with F.T. also assisting in data curation. All authors read the manuscript and agreed to publish the current version.

## Notes

### Competing Interest Statement

The authors have declared no competing interest.

### Summary of Updates

In the original version, the order of the article's authors is incorrect; the author order has been corrected.

